# Hepcidin and conventional markers to detect iron deficiency in severely anaemic HIV-infected patients in Malawi

**DOI:** 10.1101/666727

**Authors:** Minke HW Huibers, Job C Calis, Theresa J Allain, Sarah E. Coupland, Chimota Phiri, Kamija S Phiri, Dorien W Swinkels, Michael Boele van Hensbroek, Imelda Bates

**Affiliations:** Global child health group, Emma Children’s Hospital, Amsterdam University Medical Centres, University of Amsterdam, The Netherlands; Amsterdam Institute of Global Health Development (AIGHD), Amsterdam, the Netherlands; Department of Pediatric Intensive Care, Emma Children’s Hospital, Amsterdam University Medical Centres, University of Amsterdam, the Netherlands; Department of Paediatrics, College of Medicine, Queen Elizabeth Central Hospital, Blantyre, Malawi; Liverpool School of Tropical Medicine, Liverpool, United Kingdom; Department of Internal Medicine, College of Medicine, Queen Elizabeth Central Hospital, Blantyre, Malawi; Department of Molecular and Clinical Cancer Medicine, Pathology, University of Liverpool, Liverpool, United Kingdom; Department of Pathology, Liverpool Clinical Laboratories, Royal Liverpool University Hospital, Liverpool, UK; School of Public Health and Family Medicine, College of Medicine, Blantyre, Malawi; Department of Laboratory Medicine, Radboud university medical centre, Nijmegen, the Netherlands; Hepcidinanalysis.com, Nijmegen, the Netherlands

## Abstract

**Introduction:** Iron deficiency is a treatable cause of severe anaemia in low-and-middle-income-countries (LMIC). Diagnosing it remains challenging as peripheral blood markers poorly reflect bone-marrow iron deficiency (BM-ID), especially in the context of HIV-infection.

**Methods:** Severe anaemic (haemoglobin ≤70g/l) HIV-infected adults were recruited at Queen Elizabeth Central Hospital, Blantyre, Malawi. BM-ID was evaluated. Accuracy of blood markers including hepcidin alongside mean corpuscular volume, mean cellular haemoglobin concentration, serum iron, serum ferritin, soluble transferrin receptor (sTfR), sTfR -index, sTfR–ratio to detect BM-ID was valued by ROC area under the curve (AUC^ROC^).

**Results:** Seventy-three patients were enrolled and 35 (48.0%) had BM-ID. Hepcidin and MCV performed best; AUC^ROC^ of 0.593 and 0.545. Other markers performed poorly (ROC<0.5). The AUC^ROC^ of hepcidin in males was 0.767 (sensitivity 80%, specificity 78%) and in women 0.490 (sensitivity 60%, specificity 61%).

**Conclusion:** BM-ID deficiency was common in severely anaemic HIV-infected patients and is an important and potential treatable contributor to severe anaemia. Hepcidin was the best, though still suboptimal, marker of BM-ID. Hepcidin, which is directly linked to iron absorption, is a very promising marker to guide curative iron supplementation policies in severely anaemic HIV-infected patients.

## Introduction

Anaemia affects approximately a third of the world’s population and substantially reduces the disability- adjusted life years worldwide (1). Iron deficiency contributes to development of anaemia and is diagnosed in more than half of all anaemic persons (2). Consequently, iron supplements remain the backbone of prevention and treatment protocols for anaemia.

Anaemia has an extensive list of potential causes. In sub-Saharan Africa, where this condition is most common, its aetiology is even more complex and in these setting aetiologies commonly co-incide requiring a multifactorial approach (3, 4). HIV may be the cause of anaemia by its direct effect on BM cells, but can also increase the range of aetiological factors to encompass opportunistic viral, bacterial and parasitic infections, drugs such as Zidovudine and co-trimoxazole, micronutrient deficiencies and neoplastic diseases (5, 6).

The exact role of iron deficiency, one of the few potentially preventable and treatable causes of anaemia, remains unclear due to its diagnostic challenges in HIV-infected patients in low resource settings (3, 4, 7). Peripheral blood markers, including erythrocyte indices, serum iron, ferritin, and soluble transferrin receptor (sTfR), have been evaluated but their accuracy is often negatively affected by inflammatory states and renal and liver conditions, which are common in both the African and HIV-infected populations (8–11). Previous studies therefore concluded that the uses of peripheral blood markers, such as ferritin, are not reliable without correction for inflammation (12). The evaluation of iron in the bone marrow is considered the ‘gold standard’ to diagnose iron deficiency, but bone marrow sampling is invasive and requires skilled staff for sampling and interpretation, which is challenging in low resource settings. Moreover, for large-scale use, a reliable peripheral blood marker is needed to replace bone-marrow biopsy to predict bone marrow iron deficiency (BM-ID).

Hepcidin is a relatively new marker, which regulates iron absorption from the gastrointestinal tract and iron release from stores, both of which are important pathways controlling the availability of iron for incorporation in the erythrocyte precursors (13). Increases of iron plasma levels stimulate the production of hepcidin, which blocks further iron absorption from the gastrointestinal tract and iron release from storage. However, hepcidin is also an acute phase protein and serum levels increases during infections (14).

We investigated the prevalence of BM-ID in HIV-infected Malawian adult patients with severe anaemia. We further evaluated the accuracy of peripheral blood markers, as well of hepcidin to identify BM-ID in this population.

## Methods

From February 2010 to March 2011, all adults admitted to the Department of Internal Medicine of the Queen Elizabeth Central Hospital (QECH), Blantyre, Malawi with a diagnosis of severe anaemia and HIV infection were approached for informed consent and study enrolment. This study is a sub-study of the larger observational cohort study (n=199) concerning severely anaemic (haemoglobin ≤ 70 g/l) HIV-infected patients. Bone marrow sampling was performed if the patient consented and the patient was clinically stable. Of the 199 included patients, 73 BM (37%) samples were included in this sub-study as the BM sampling was performed and the quality of the sample taken was appropriate.

### Methods| Laboratory assays and blood markers

Haemoglobin concentration was measured on admission using the HemoCue B-Haemoglobin analyser (HemoCue, Ängelholm, Sweden) to screen patients for eligibility. After informed consent a venous blood sample was collected and bone marrow sample taken from the iliac crest. All blood samples were analysed within 24 hours of collection or stored at −80°C. Haemoglobin and red cell indices (MCV, MCH and MCHC) were determined using an automated haematology analyser (Beckman Coulter, Durban, South Africa). CD4-cell counts were assessed using BD FACS Count (BD Biosciences, San Jose, CA, USA). Transferrin, iron, ferritin, folate and vitamin B12 were analysed on Modular P800 and Monular Analytic E170 systems (Roche Diagnostics, Switzerland). Soluble transferrin receptor (sTfR) levels were measured using ELISA (Ramco Laboratories, TX, USA). Commonly used ratios to define iron deficiency were calculated including the sTfR-index: sTfR (mg/L) divided by log ferritin (ug/L); and the ‘sTfR ratio’: sTfR (mg/L) x1000/ ferritin (ug/L) level (15). International accepted cut-offs were applied. For sTfR we used 2.75 mg/l and 3.6 mg/l and for the sTfR-index: 1.8 and 2.2, 2.8 respectively, as no international cut-offs have been defined, these represented the most resent consensus (16).

Serum hepcidin-25 measurements were performed in December 2012/January 2013 (Testing lab: Hepcidinanalysis.com, Nijmegen, The Netherlands) by a combination of weak cation exchange chromatography and time-of-flight mass spectrometry (WCX-TOF MS) using synthetic hepcidin-24 as internal standard (17–19). Peptide spectra were generated on a Microflex LT matrix-enhanced laser desorption/ionisation TOF MS platform (Bruker Daltonics, Bremen, Germany). Hepcidin concentrations are expressed as nanomoles per litre (nmol/L). The lower limit of detection of this method was 0.5 nmol/L (18).

### Methods| BM-ID

Aspirate samples were spread onto slides and trephine biopsies were fixed, decalcified and embedded in paraffin wax (20, 21). Bone marrow samples were sent to the Haematology- Pathology Referral Centre at the Royal Liverpool University Hospital, Liverpool UK, for analysis. Sections of the trephine blocks were stained with Perls’ Prussian Blue stain to detect iron stores(21). Intracellular iron in bone marrow trephine blocks was graded using the Stuart-Smith scale, which classifies the iron content of bone marrow into six grades (0–6). For bone marrow smears, iron was graded using Gale’s grading (0-4). Iron deficiency was defined as no visible or severe reduced iron particles in a few reticulum cells under high power magnification; grade 0-1 on both scales (22, 23).

### Methods| Infections

HIV infection was confirmed using two point-of-care antibody tests (Unigold^®^ and Determine^®^). Different types and severity of on-going infections were evaluated. Including; HIV: CD4 counts ≤200 cells/mm^3^ and/or viral load >1000 copies/ml. Malaria: presence of malaria parasites in a thick blood film assessed by light microscopy. Tuberculosis (TB) defined as one or more of the following: a) positive sputum culture; b) chest X-ray with signs of pulmonary tuberculosis and/or; c) on-going TB treatment at time of enrolment; d) clinical diagnosis based on generalized lymphadenopathy and/or night sweats > 30 days with unknown origin; e) caseating granulomata in the bone marrow trephine. Bacteraemia was defined as blood cultures growing potential pathogen including streptococcus, enterococcus and micrococcus species, non-Typhoid Salmonella and Klebsiella pneumonia. Furthermore, viral infections including Parvo-B19, Cytomegalovirus (CMV) and Epstein-Barr virus (EBV) were evaluated by PCR and defined as positive by viral load >100 copies/ml.

### Methods| Ethics

The Research Ethics Committee of the College of Medicine, University of Malawi (P.09.09.824) and the Research Ethics Committee of Liverpool School of Tropical Medicine (research protocol 09.64) approved the study. The purpose of the study was explained to the patients in the local language (Chichewa), and written informed consent was obtained before inclusion into the study.

### Methods| Statistics

The data were analysed using Stata (version 12) (STATA Corp. LP, Texas, TX, USA). Baseline characteristics were compared between BM-ID and non-deficient patients using Chi-square test (dichotomous data) or t-test (continuous) or Pearson Chi-square test (continuous not normally distributed). Confounding was enhanced to evaluate hepcidin concentrations, gender, HIV disease progression, the use of ART as baseline, and TB infection (Pearson Chi-square test). The p-values reported are two-sided, and a level of p<0.05 was interpreted as significant.

The accuracy of the different peripheral blood markers, including hepcidin, to discriminate BM-ID were evaluated by receiver operating characteristics curves (ROC)(24). Corresponding areas under the curve (AUC^ROC^) were created. AUC^ROC^ measures the two-dimensional area underneath the ROC curve and provides a summative measure of performance across all possible classification thresholds (25). The AUC^ROC^ <0.70 is weighed as low diagnostic; AUC^ROC^ of 0.70–0.90 as moderate diagnostic and a AUC^ROC^ ≥0.90, high diagnostic accuracy (26). Sensitivity and specificity were calculated for predefined internationally accepted cut-offs (8, 15, 16, 27). For hepcidin the best cut-off value for diagnosing BM-ID were determined using ROC-curve analyses with the Youden index (maximum (sensitivity + specificity – 1))(25). As gender differences are known for hepcidin (30) hepcidin outcome was hence evaluated by gender.

## Results

Of the 73 HIV-infected adults in our sub-study, a total of 45 (61.6%) had severe anaemia (Hb 50-<70g/dL) and 28 (38.4 %) had very severe anaemia (Hb<50g/dL). The mean patient age was 33.7 (SD 8.7) years, and 43 (58.9%) patients were female. A CD4 count ≤ 200 cells/mm^3^ was seen in 31/56 (55.4%) and a viral load >1000 copies/ml was present in 57/76 (75.0%) of patients. A total of 34/73 (46.6%) patients had been started on anti-retroviral ART treatment, of which most were on first line treatment at time of the study (Efavirenz, Lamivudine, Tenofovir). The most common infections in this population were tuberculosis (39/73; 53.4%) and EBV (30/45; 66.7%). All baseline characteristics are shown in table 1.

**Table 1.** Baseline characteristics in this population of severely anaemic HIV patients’ stratified according to bone marrow iron deficiency (BM-ID). Abbreviations: ART: antiretroviral therapy. BMI: Body mass index. TB: Tuberculosis. ^1^ First line ART include combination of Stavudine (d4T), Lamivudine (3Tc) and Nevirapine (NVP) (28). ^2^ Advanced HIV disease including a CD4 count ≤ 200 cells/mm^3^ and/or viral load > 1000 copies/ml. ^3^Bacteraemia; a blood culture with clean growing potential pathogen including streptococcus (41.7%; 5/12), enterococcus (16.7%;2/12) and non-Typhoid Salmonella (16.7%;2/12). ^4^Malaria: presence of malaria parasites on a thick blood film. ^5^Tuberculosis (TB): one or more of the following present: a) positive sputum culture, b) chest X-ray with signs of pulmonary tuberculosis and/or c) on-going TB treatment at time of enrolment d) clinical diagnosis by local doctor including unknown generalized lymphadenopathy and/or night sweats > 30 days with unknown origin e) caseating granulomata in the bone marrow trephine. ^6^ Epstein-Barr, cytomegalo- and parvo-B19 virus infection are diagnosed by a virus load of 1000 copies/ml. Abbreviations: MCV; mean cellular volume, MCH; mean corpuscular haemoglobin, s-TfR: Soluble transferrin receptor, TfR-index (sTfR(mg/L) /Log ferritin(ug/L)), TfR Ratio (sTrR(mg/L))x1000/ferritin(ug/L)).

| Characteristic | Overall | Non BM-ID | BM-ID | P-value |
| --- | --- | --- | --- | --- |
| **BM-ID** | **35/73 (48.0%)** | **38/73 (52.1%)** | **35/73 (48.0%)** |  |
| Age, years (mean, SD) | 33.7 (8.7) | 32.7 (8.6) | 34.7 (8.9) | 0.331 |
| Gender (female) (%) | 43/73 (58.9%) | 19/38 (50.0%) | 24/35 (68.6%) | 0.107 |
| **Haematology & iron markers** |  |  |  |  |
| Very severe anaemia (Hb≤ 50g/l) (%) | 28/73 (38.4%) | 13/38 (34.2%) | 15/35 (42.9%) | 0.448 |
| Haemoglobin (Hb)(g/l), (median, IQR) | 56.0 (43.0-63.0) | 58.5 (45.0-64.0) | 54.0 (36.0-63.0) | 0.136 |
| MCV (fl), (median, IQR) | 85.8 (79.4-98.1) | 87.3 (79.6-99.0) | 83.5 (79.1-94.7) | 0.534 |
| MCH (pg/cells), (mean, SD) | 29.0 (5.9) | 23.6 (6.1) | 26.3 (5.5) | 0.084 |
| Serum iron (umol/l), (median, IQR) | 5.1 (3.3-11.1) | 4.7 (3.0-7.9) | 5.6 (3.8-22.2) | 0.053 |
| Ferritin (ug/dL), (median, IQR) | 87.2 (49.6-100.0) | 87.1 (50.1-97.1) | 87.9 (36.0-100.0) | 0.488 |
| sTfR receptor (mg/l), (median, IQR) | 2.9 (1.6-3.7) | 2.8 (1.7-3.7) | 3.0 (1.2-3.9) | 0.934 |
| sTfR index (median, IQR) | 1.6 (0.8-2.2) | 1.5 (0.9-2.1) | 1.6 (0.6-2.6) | 0.901 |
| sTfR Ratio (median, IQR) | 35.1 (17.5-69.3) | 33.2 (21.3-61.9) | 36.6 (11.5-79.6) | 0.747 |
| Hepcidin (ng/ml) (median, IQR) | 7.3 (3.3-13.3) | 9.2 (4.9-13.2) | 5.1 (3.1-13.7) | 0.196 |
| **HIV disease and treatment** |  |  |  |  |
| ART at enrolment (%) | 34/73 (46.6%) | 20/38 (52.6%) | 14/35 (40.0%) | 0.280 |
| CD4 count | 31/56 (55.4%) | 15/28 (53.6%) | 14/25(56.0%) | 0.859 |
| ≤ 200 cells/mm<sup>3</sup> |  |  |  |  |
| Viral load<br>>1000 copies/ml | 57/76 (75.0%) | 25/38 (65.8%) | 30/35 (85.7%) | 0.048 |
| **Infection(s)** |  |  |  |  |
| Bacteraemia<sup>3</sup> | 12/73 (16.4%) | 6/38 (15.8%) | 6/35 (17.1%) | 0.876 |
| Malaria<sup>4</sup> | 3/63 (4.7%) | 2/32 (6.3%) | 1/31 (3.2%) | 0.573 |
| Tuberculosis<sup>5</sup> | 39/73 (53.4%) | 20/38 (52.6%) | 19/35 (54.3%) | 0.877 |
| Epstein-Barr virus<sup>6</sup> | 30/45 (66.7%) | 19/26 (73.1%) | 11/19 (57.9%) | 0.286 |
| Cytomegalo virus<sup>6</sup> | 18/54 (33.3%) | 11/28 (39.3%) | 7/26 (26.9%) | 0.336 |
| Parvo-B19 virus<sup>6</sup> | 1/59 (1.7%) | 0/27 (-) | 1/32 (3.1%) | 0.230 |
| **Nutritional status** |  |  |  |  |
| Underweight<br>(BMI < 18.5) (%) | 22/49<br>(44.9%) | 8/24<br>(33.3%) | 14/25<br>(56.0%) | 0.111 |

### Results| BMI-ID and blood markers

BM-ID was seen among 35 (48.0%) of the patients, table 1. The performances of the peripheral blood markers to diagnose BM-ID are displayed in table 2. All markers displayed low diagnostic accuracy (AUC^ROC^< 0.7). MCV had the highest AUC^ROC^ value of the common peripheral blood markers (0.545), the sensitivity and specificity using the common cut off of 83fL were 42% and respectively 67%. The use of hepcidin to detect BM-ID resulted in an AUC^ROC^ 0.593. We stratified the analysis for hepcidin according to gender; the AUC^ROC^ for men and women was 0.767 and 0.490 respectively. The optimal hepcidin concentration for the detection of BM-ID was ≤7 ng/ml; sensitivity 67% & specificity 67%. In males the optimum cut off was ≤6 ng/ml (sensitivity 80%; specificity 78%); whilst for women this was ≤7 ng/ml (sensitivity 60%; specificity 61%, figure 1). The hepcidin concentration did not differ significantly by gender (p=0.831), HIV disease progression (p=0.819), the use of ART at enrolment (p=0.616), and TB infection (p=0.590) in a univariate analysis.

**Table 2.** Accuracy of peripheral blood markers to detect bone marrow iron deficiency (gold standard). Abbreviations: AUC: area under curve of receiver operating characteristic (ROC), where 0.5 would be expected by chance and 1 denotes a perfect test. 95%-CI: 95% confidence interval. MCV; mean cellular volume, MCH; mean corpuscular haemoglobin, sTfR: Soluble transferrin receptor, sTfR-index (sTfR(mg/L) /Log ferritin(ug/L)), sTfR Ratio (sTrR(mg/L))x1000/ferritin(ug/L)).^1^ (29) ^2^ (11) ^3 (29) 4(15, 16)^

| Potential markers | AUC<sup>ROC</sup> | 95%-CI | Cut-off | Sensitivity | 95%-CI | Specificity | 95%-CI |
| --- | --- | --- | --- | --- | --- | --- | --- |
| MCV (fl)<sup>1</sup> | 0.545 | 0.404-0.685 | ≤83 | 42% | 25.2-58.8% | 67% | 51.0-83.0% |
| MCH (pg/cells) <sup>1</sup> | 0.365 | 0.230-0.499 | ≤27 | 52% | 35.2-69.0% | 29% | 13.7-44.3% |
| Serum iron (μmol/l)<br><sup>1</sup> | 0.368 | 0.239-0.498 | ≤ 10 | 60% | 43.8-76.2% | 18% | 5.8-30.2% |
| Ferritin (μg/l) <sup>1</sup> | 0.441 | 0.293-0.588 | ≤30 | 13% | 1.0-25.0% | 88% | 76.7-99.3% |
|  |  |  | ≤70 | 30% | 13.6-46.4% | 66% | 49.6-82.4% |
| sTfR receptor<br>(mg/l)<sup>3</sup> | 0.522 | 0.378-0.667 | ≥1.8 | 71% | 55.0-87.0% | 29% | 13.7-44.3% |
|  |  |  | ≥2.8 | 58% | 40.6-73.1% | 56% | 39.3-72.7% |
|  |  |  | ≥3.6 | 32% | 15.6-48.2% | 76% | 61.6-99.4% |
|  |  |  | ≥8.0 | 3% | 0-9.0% | 100% | - |
| sTfR index <sup>4</sup> | 0.523 | 0.375-0.672 | ≥1.8 | 47% | 29.1-64.9% | 67% | 46.3-79.7% |
|  |  |  | ≥2.2 | 27% | 11.1-42.9% | 75% | 60.0-90.0% |
|  |  |  | ≥2.8 | 20% | 5.7-34.3% | 81% | 67.4-94.6% |
|  |  |  | ≥3.5 | 10% | 0-20.7% | 97% | 75.1-86.9% |
| sTfR Ratio <sup>4</sup> | 0.508 | 0.359-0.656 | ≥100 | 17% | 0-23.4% | 91% | 71.1-90.9% |

**Figure 1.**
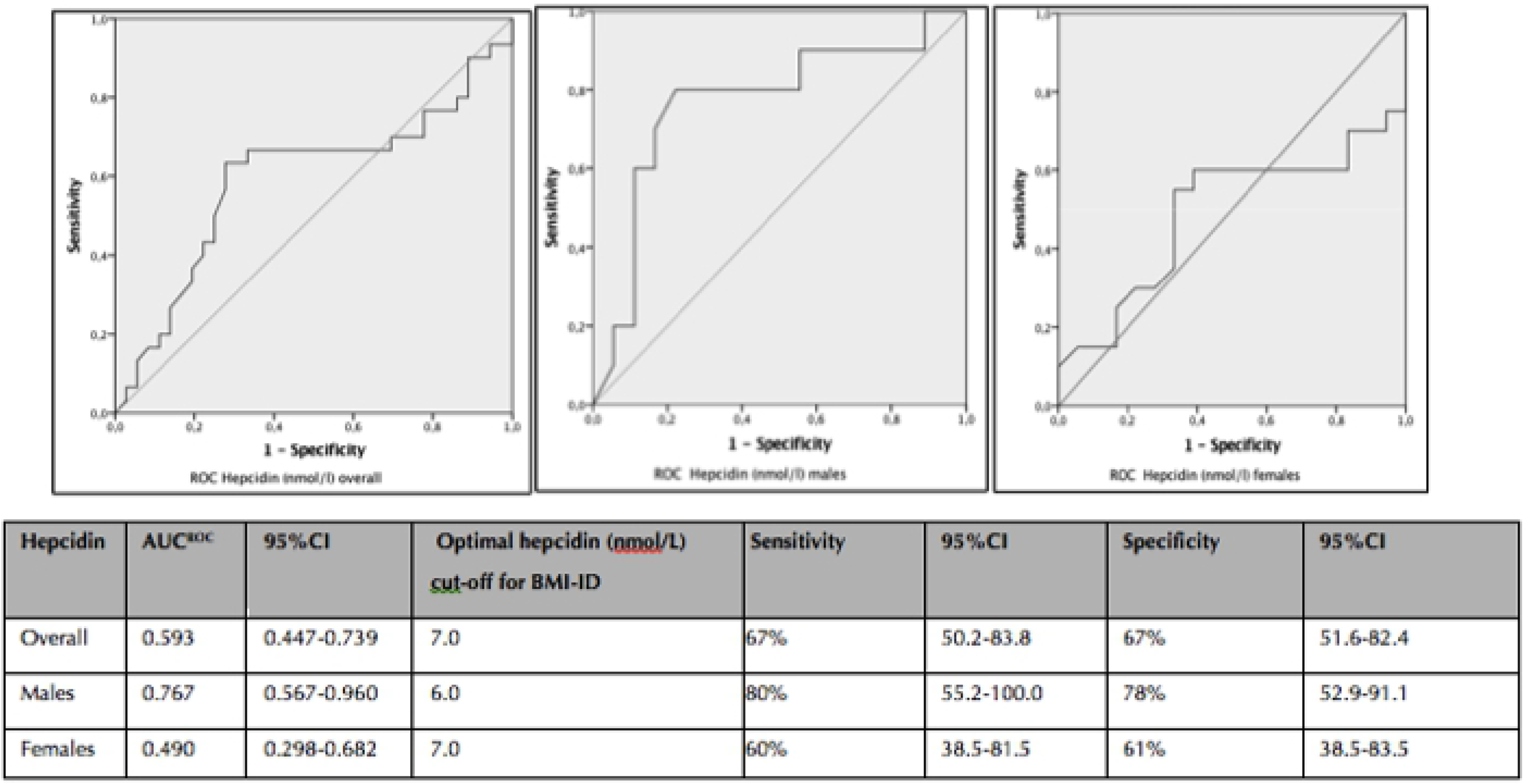
Hepcidin (nmol/L) ROC curve by gender with optimal cut-off. The best cut-off value for diagnosing BM-ID was determined by the Youden index (maximum (sensitivity + specificity −1)) in the ROC-curve (25). Abbreviations: AUC^ROC^: area under curve of receiver operating characteristic.

## Discussion

In this study on hepcidin and conventional markers to detect BM-ID in severely anaemic HIV-infected patients in Malawi, we found that BM-ID was present in almost half of our patients. In the first study evaluating hepcidin as a marker for BM-ID among severely anaemic HIV-infected adults in such a setting we found that hepcidin had the highest AUC^ROC^ 0.593 amongst all peripheral markers and therefore was the best marker to use for detecting BM-ID. MCV was found to be the best conventional peripheral blood marker for BM-ID. As the MCV is commonly provided as part of routine full blood counts this marker may be of some use in resource-limited settings.

BM-ID was highly prevalent among our population of HIV-infected and severely anaemic patients. The prevalence of BM-ID in this study is higher than previous reports on similar populations that were published in the pre-ART era when BM-ID was reported to be 18%-25% in severely anaemic HIV patients (8, 30). The increased prevalence of BM-ID may be explained by the effect of ART. Previous reports on HIV-associated anaemia before 2010 included patients who were mostly ART naïve and advanced HIV disease and/or severe immune suppression were common (8). Although VL above 1000 copies/ml (75%) and CD4 counts below 200 cells/ml were (55.4%) still common in the presenting population, median CD4 counts (325 cells/mm^3^) were much higher than in the previous reports (median 67 cells/ml) (4, 8). Initiation of ART aims to stop HIV disease progression, promote immune reconstitution and reduce the risk of (opportunistic) diseases. This may also be reflected in our cohort, and has likely changed the aetiology of severe anaemia among HIV-infected patients as compared to older studies. In this group of HIV-infected patients with better immune systems the aetiology of severely anaemic may be more similar to the aetiology of non-HIV infected patients. The BM-ID prevalence of 48% is comparable to previous findings among HIV-uninfected African populations with severe anaemia, which supports this hypothesis (8). Our data need to be confirmed in the on-going ‘treat all’ era but they could have great impact on preventive and curative policies concerning iron supplementation in severely anaemic HIV-infected patients. Irrespective of the cause, the role of iron supplementation to prevent and treat severe anaemia appears to have gained importance.

Our results concerning the accuracy of peripheral blood markers to detect BM iron deficiency indicate that it is not easy to reliably detect those with deficient BM iron stores, which corroborates with previous studies (8, 9). Hepcidin, a specific hormone in metabolising iron, did perform slightly better than conventional markers, but remained far from good or even perfect. Additionally, hepcidin is a key player in the absorption of iron and thus may be used to not only select those needing iron but also may predict iron supplementation, response, safety and timing. Hepcidin as a possible marker for BM-ID, has not been evaluated before in this population of severely anaemic HIV-infected adults in Africa. It is not surprising that hepcidin remains from prefect as a marker for BM-ID as we know that hepcidin levels are also affected by inflammation which is highly present among HIV-infected patients, especially living in resource-limited settings (31–33). For example, when hepcidin levels are high, the absorption of dietary iron and release of macrophage iron to serum are blocked as protection, resulting in a relative hypoferremia and an increase iron into the macrophages, which is thought to be anti-infective. Consequently during malaria or TB infection or immune deficiency with low CD4 counts, hepcidin levels are increased (32, 33). Among children in Malawi, including children with HIV, our group previously reported low hepcidin levels (34). Further, hepcidin was suggested as a possible useful marker in guiding iron therapy in severely anaemic children, as low hepcidin levels were related toward a diminished expected up regulation of hepcidin by inflammation and iron deficiency due to an increase of erythropoietin in this population (34). Our data on hepcidin and (standardized) identified cut-offs are highly likely to be relevant as there is a need for a reliable marker to define BM-ID and to start iron supplementation among HIV-infected patients in resource limited settings such as Malawi. Intervention studies, using hepcidin as a marker, should be performed to assess feasibility and effect of such an intervention.

Worldwide hepcidin concentrations are measured by various methods, which differ considerably in absolute hepcidin concentrations (35). Recently, secondary hepcidin reference material, that has been value assigned by a primary reference material, has become available (36). Standardization in February 2019 of a similar hepcidin assay as we used in 2012/2013 for our study, resulted in (only) 5.4 % increase of hepcidin concentrations (C. Laarakker and D. Swinkels unpublished data) (37). For this specific patient population, our study thus provides a first and rough estimate for cut off point that are universally applicable by other assays that they are standardized using this same reference material. However, for formal universal use of these cut-off points these values should be confirmed by studies that directly measure samples with a standardized hepcidin method. Additionally, hepcidin optimal cut-offs were more sensitive and lower for man in comparison to woman. A direct clarification for this effect is challenging. Age and gender differences in hepcidin concentrations are known. Within the mean age group of our study population, 30-35 years of age, (normal) hepcidin concentrations were reported to be higher on man than woman (36, 38). However reference data available is coming from Europe and no hepcidin reference levels are known for an African population. Therefore a direct comparison is challenging. At last an explanation(s) can be found in levels of infection or control of the HIV disease, which in our population was not different for woman and man (data not shown). Outcome should be formally confirmed with studies that directly measure with the standardized hepcidin method to enhance confirmed hepcidin cut-offs in this specific patient population.

Iron deficiency is treatable and preventable, however supplementation has been associated with an increased number of severe infections, including an increase in malaria, which, especially in an immune compromised population, may be dangerous (39, 40). Therefore a reliable diagnosis of iron deficiency is important, as supplementation will put the patient at risk. Our study underlined that the current used and known peripheral blood markers performed poorly. For several decades clinicians and researchers have evaluated peripheral markers to diagnose BM-ID in laboratory resource limiting settings like Malawi and reported poor performance which is commonly considered to be caused by inflammatory conditions which are common in African and especially in HIV infected patients (8, 9). Some of the markers tested, such as sTfR concentrations, alone or in combination with other markers, were designed to better reflect iron stores (41) irrespective of inflammation (42). These markers did not perform well in this study either. Potential explanations for the poor performances of sTfR in our population may be the lack of clear cut-offs, as suggested by other studies (8) and the fact that sTfR is also influenced by erythropoietin, which may play an (even more) important role in severe HIV-associated anaemia (32). The best performing conventional peripheral blood marker was MCV. Microcytosis is commonly used as a screening test for deficiency (27, 43); however, MCV was never found to be an accurate predictor of BM-ID (8, 9, 44). Although MCV did not have a high sensitivity or specificity and the AUC^ROC^ was of low diagnostic value, it remains of some use in this population as it was the best available common marker and it is relatively easily available as part of automated full blood counts.

Our study has several shortcomings. Firstly, bone marrow testing was only performed in a subset of patients, which may have introduced a sampling bias. Reasons for not taking bone marrow included a severe clinical condition of the patient or patients not consenting to this aspect of the study. However, it is one of the largest studies with BM results to date. Secondly, our study was performed in 2010 when antiretroviral treatment (ART) was provided according to the national and hospital guidelines. Accordingly, ART could only be started in the outpatient ART clinic after discharge from hospital. Currently ART is started much earlier in the course of HIV infection so our study patients are likely to have had more advance disease than current patients. Nevertheless, this is the first study combining bone marrow data with a large set of peripheral blood markers, including hepcidin, in a group of HIV-infected severely anaemic African patients. We believe our study provides important information, which is valuable for clinicians who care for HIV-infected persons with severe anaemia in Malawi and other resource-limited settings.

## Conclusion

Bone marrow iron deficiency was present in almost half of severely anaemic HIV-infected adults. This, substantial increase compared to data from the pre-ART era, underline the potential importance of preventive and therapeutic role of iron supplementation to reduce the problem of severe anaemia in HIV-infected patients. Detection and safe treatment of BM-ID is hampered by a lack of peripheral iron markers. Hepcidin was found to be the most accurate marker and could be used to guide and predict the effect of iron supplementation. Although hepcidin evaluations are not routinely available in settings such as Malawi, our study findings are important as hepcidin could guide and predict the effect of safe iron supplementation. This is important because of the potential risk of increased infection risk due to iron supplementation, its effect and safety should be evaluated in future intervention studies.

## Acknowledgement

The authors would like to thank all of the study participants, doctors, nurses and support staff of Queens Elizabeth Hospital and the Malawi-Liverpool-Wellcome centre in Blantyre for their participation and cooperation. This study was supported by the Nutricia research foundation (Project number 2017-43), The Hague, the Netherlands and the Wellcome Trust (Project number WT086559), Liverpool, United Kingdom. The funders had no role in the study design, data collection and analysis, decision to publish or preparation of the manuscript

## References

1. Kassebaum NJ, Wang H, Lopez AD, Murray CJ, Lozano R. Maternal mortality estimates - Authors’ reply. Lancet (London, England). 2014;384(9961):2211–2.

2. Lopez A, Cacoub P, Macdougall IC, Peyrin-Biroulet L. Iron deficiency anaemia. Lancet (London, England). 2016;387(10021):907–16.

3. Calis JC, Phiri KS, Faragher EB, Brabin BJ, Bates I, Cuevas LE, et al. Severe anemia in Malawian children. The New England journal of medicine. 2008;358(9):888–99.

4. Lewis DK, Whitty CJ, Walsh AL, Epino H, Broek NR, Letsky EA, et al. Treatable factors associated with severe anaemia in adults admitted to medical wards in Blantyre, Malawi, an area of high HIV seroprevalence. Transactions of the Royal Society of Tropical Medicine and Hygiene. 2005;99(8):561–7.

5. Butensky E, Kennedy CM, Lee MM, Harmatz P, Miaskowski C. Potential mechanisms for altered iron metabolism in human immunodeficiency virus disease. The Journal of the Association of Nurses in AIDS Care : JANAC. 2004;15(6):31–45.

6. Klassen MK, Lewin-Smith M, Frankel SS, Nelson AM. Pathology of human immunodeficiency virus infection: noninfectious conditions. Annals of diagnostic pathology. 1997;1(1):57–64.

7. Marti-Carvajal AJ, Sola I, Pena-Marti GE, Comunian-Carrasco G. Treatment for anemia in people with AIDS. The Cochrane database of systematic reviews. 2011(10):Cd004776.

8. Lewis DK, Whitty CJ, Epino H, Letsky EA, Mukiibi JM, van den Broek NR. Interpreting tests for iron deficiency among adults in a high HIV prevalence African setting: routine tests may lead to misdiagnosis. Transactions of the Royal Society of Tropical Medicine and Hygiene. 2007;101(6):613–7.

9. Jonker FA, Boele van Hensbroek M, Leenstra T, Vet RJ, Brabin BJ, Maseko N, et al. Conventional and novel peripheral blood iron markers compared against bone marrow in Malawian children. Journal of clinical pathology. 2014;67(8):717–23.

10. Odunukwe NN, Salako LA, Okany C, Ibrahim MM. Serum ferritin and other haematological measurements in apparently healthy adults with malaria parasitaemia in Lagos, Nigeria. Tropical medicine & international health : TM & IH. 2000;5(8):582–6.

11. Aguilar R, Moraleda C, Quinto L, Renom M, Mussacate L, Macete E, et al. Challenges in the diagnosis of iron deficiency in children exposed to high prevalence of infections. PloS one. 2012;7(11):e50584.

12. Nel E, Kruger HS, Baumgartner J, Faber M, Smuts CM. Differential ferritin interpretation methods that adjust for inflammation yield discrepant iron deficiency prevalence. Maternal & child nutrition. 2015;11 Suppl 4:221–8.

13. Hugman A. Hepcidin: an important new regulator of iron homeostasis. Clin Lab Haematol. 2006;28(2):75–83.

14. D’Angelo G. Role of hepcidin in the pathophysiology and diagnosis of anemia. Blood research. 2013;48(1):10–5.

15. Skikne BS, Punnonen K, Caldron PH, Bennett MT, Rehu M, Gasior GH, et al. Improved differential diagnosis of anemia of chronic disease and iron deficiency anemia: a prospective multicenter evaluation of soluble transferrin receptor and the sTfR/log ferritin index. American journal of hematology. 2011;86(11):923–7.

16. Suominen P, Punnonen K, Rajamaki A, Irjala K. Evaluation of new immunoenzymometric assay for measuring soluble transferrin receptor to detect iron deficiency in anemic patients. Clinical chemistry. 1997;43(9):1641–6.

17. Swinkels DW, Girelli D, Laarakkers C, Kroot J, Campostrini N, Kemna EH, et al. Advances in quantitative hepcidin measurements by time-of-flight mass spectrometry. PloS one. 2008;3(7):e2706.

18. Kroot JJ, Laarakkers CM, Geurts-Moespot AJ, Grebenchtchikov N, Pickkers P, van Ede AE, et al. Immunochemical and mass-spectrometry-based serum hepcidin assays for iron metabolism disorders. Clin Chem. 2010;56(10):1570–9.

19. Kroot JJ, Tjalsma H, Fleming RE, Swinkels DW. Hepcidin in human iron disorders: diagnostic implications. Clin Chem. 2011;57(12):1650–69.

20. Bain BJ. Bone marrow aspiration. Journal of clinical pathology. 2001;54(9):657–63.

21. Bain BJ. Bone marrow trephine biopsy. Journal of clinical pathology. 2001;54(10):737–42.

22. Gale E, Torrance J, Bothwell T. The quantitative estimation of total iron stores in human bone marrow. The Journal of clinical investigation. 1963;42:1076–82.

23. Finch CA. The detection of iron overload. The New England journal of medicine. 1982;307(27):1702–4.

24. Metz CE. Basic principles of ROC analysis. Seminars in nuclear medicine. 1978;8(4):283–98.

25. Bewick V, Cheek L, Ball J. Statistics review 13: receiver operating characteristic curves. Critical care (London, England). 2004;8(6):508–12.

26. JA. S. Signal Detection Theory and ROC Analysis in Psychology and Diagnostics Collected Papers: Mahwah, Erlbaum; 1996. p. 94–117.

27. Bates I. Practical hematology. Chapter 2: reference ranges and normal values. 2018.

28. Ministry of Health M. Treatment of AIDS, guidelines for the use of antiretroviral treatment in Malawi,edition 2010. 2010(Third edition since 2008).

29. Rimon E, Levy S, Sapir A, Gelzer G, Peled R, Ergas D, et al. Diagnosis of iron deficiency anemia in the elderly by transferrin receptor-ferritin index. Archives of internal medicine. 2002;162(4):445–9.

30. Kagu MB, Khalil MI, Ahmed SG. Bone marrow macrophage iron stores in patients with HIV infection and AIDS-associated Kaposi’s sarcoma. African journal of medicine and medical sciences. 2007;36(2):125–8.

31. Prentice AM, Doherty CP, Abrams SA, Cox SE, Atkinson SH, Verhoef H, et al. Hepcidin is the major predictor of erythrocyte iron incorporation in anemic African children. Blood. 2012;119(8):1922–8.

32. Wisaksana R, de Mast Q, Alisjahbana B, Jusuf H, Sudjana P, Indrati AR, et al. Inverse relationship of serum hepcidin levels with CD4 cell counts in HIV-infected patients selected from an Indonesian prospective cohort study. PloS one. 2013;8(11):e79904.

33. de Mast Q, Syafruddin D, Keijmel S, Riekerink TO, Deky O, Asih PB, et al. Increased serum hepcidin and alterations in blood iron parameters associated with asymptomatic P. falciparum and P. vivax malaria. Haematologica. 2010;95(7):1068–74.

34. Jonker FA, Calis JC, Phiri K, Kraaijenhagen RJ, Brabin BJ, Faragher B, et al. Low hepcidin levels in severely anemic malawian children with high incidence of infectious diseases and bone marrow iron deficiency. PloS one. 2013;8(12):e78964.

35. van der Vorm LN, Hendriks JC, Laarakkers CM, Klaver S, Armitage AE, Bamberg A, et al. Toward Worldwide Hepcidin Assay Harmonization: Identification of a Commutable Secondary Reference Material. Clin Chem. 2016;62(7):993–1001.

36. Diepeveen LE, Laarakkers CMM, Martos G, Pawlak ME, Uguz FF, Verberne K, et al. Provisional standardization of hepcidin assays: creating a traceability chain with a primary reference material, candidate reference method and a commutable secondary reference material. Clinical chemistry and laboratory medicine. 2018.

37. Refference values hepcidin: www.hepcidineanalysis.com/provided-service/reference-values; 2019 [

38. Galesloot TE, Vermeulen SH, Geurts-Moespot AJ, Klaver SM, Kroot JJ, van Tienoven D, et al. Serum hepcidin: reference ranges and biochemical correlates in the general population. Blood. 2011;117(25):e218–25.

39. Esan MO, van Hensbroek MB, Nkhoma E, Musicha C, White SA, Ter Kuile FO, et al. Iron supplementation in HIV-infected Malawian children with anemia: a double-blind, randomized, controlled trial. Clinical infectious diseases : an official publication of the Infectious Diseases Society of America. 2013;57(11):1626–34.

40. Sazawal S, Black RE, Ramsan M, Chwaya HM, Stoltzfus RJ, Dutta A, et al. Effects of routine prophylactic supplementation with iron and folic acid on admission to hospital and mortality in preschool children in a high malaria transmission setting: community-based, randomised, placebo-controlled trial. Lancet (London, England). 2006;367(9505):133–43.

41. Esan MO, Jonker FA, Hensbroek MB, Calis JC, Phiri KS. Iron deficiency in children with HIV-associated anaemia: a systematic review and meta-analysis. Transactions of the Royal Society of Tropical Medicine and Hygiene. 2012;106(10):579–87.

42. Rohner F, Namaste SM, Larson LM, Addo OY, Mei Z, Suchdev PS, et al. Adjusting soluble transferrin receptor concentrations for inflammation: Biomarkers Reflecting Inflammation and Nutritional Determinants of Anemia (BRINDA) project. The American journal of clinical nutrition. 2017;106(Suppl 1):372s–82s.

43. Volberding PA, Levine AM, Dieterich D, Mildvan D, Mitsuyasu R, Saag M. Anemia in HIV infection: clinical impact and evidence-based management strategies. Clinical infectious diseases : an official publication of the Infectious Diseases Society of America. 2004;38(10):1454–63.

44. Rabindrakumar MSK, Pujitha Wickramasinghe V, Gooneratne L, Arambepola C, Senanayake H, Thoradeniya T. The role of haematological indices in predicting early iron deficiency among pregnant women in an urban area of Sri Lanka. BMC hematology. 2018;18:37.

